# Rapid, time-resolved proximity labeling by sbp1 identifies a porin domain protein at the malaria parasite periphery

**DOI:** 10.1101/2022.06.30.498261

**Authors:** David Anaguano, Carrie F. Brooks, David W. Cobb, Vasant Muralidharan

## Abstract

The deadly human malaria-causing parasite, *Plasmodium falciparum* relies on its capacity to completely remodel its host red blood cell (RBC) through the export of hundreds of parasite proteins across several membranes to the RBC. Among these exported proteins are numerous membrane proteins that are inserted into the parasite plasma membrane (PPM) during their transport via the secretory pathway. It is not known how these exported membrane proteins are extracted from the PPM for export. To answer this question, we fused the exported membrane protein skeleton binding protein 1 (SBP1) with the rapid, efficient, and promiscuous biotin ligase known as TurboID (SBP1^TbID^). Our data show that the SBP1^TbID^ fusion protein was exported efficiently to the host RBC and was able to rapidly biotinylate proteins at the host-parasite interface during its export as well as at its final destination in the host RBC. Using time-resolved proximity biotinylation and label-free quantitative proteomics, we identified early (pre-export) interactors and late (post-export) interactors of SBP1^TbID^. This led to the identification of 24 proteins that were 10-fold or more enriched in the pre-export time point compared to the post-export time point. Among these early interactors were two promising membrane-associated proteins, one of which has a predicted porin domain, that could potentially act as a translocon at the PPM for exported membrane proteins (Plasmodium translocon of exported membrane proteins or PTEM). Both proteins localize to the host-parasite interface during early stages of the intraerythrocytic cycle and conditional knockdown of these candidates show that they play essential roles in the asexual lifecycle of the parasite. Taken together, our data suggest that these two proteins may play a role in extracting membrane proteins from the PPM for export to the host RBC.

## INTRODUCTION

Malaria is a major global health issue with an estimated 241 million cases and 627 000 deaths reported during 2020^1^. This life-threatening disease is caused by apicomplexan parasites of the genus *Plasmodium*, however one species, *P. falciparum*, is the most virulent and lethal, accounting for 95% of all malaria deaths^2^. The malaria symptoms include headaches, myalgia, high fevers, severe anemia, pulmonary and renal failure, vascular obstruction, and cerebral damage. These disorders are a consequence of parasite proliferation within human red blood cells (RBC) and can persist even after parasite clearance^2,3^.

To establish infection during their intraerythrocytic cycle, *P. falciparum* parasites must extensively remodel the morphology and physiology of the RBCs. This transformation requires the export of several hundred proteins (about 10% of the parasite proteome) across the parasitophorous vacuole (PV) into the RBC cytoplasm and membrane^4–7^, and leads to increased permeability, loss of cell deformability, and formation of virulence-associated knobs at the RBC membrane^8,9^. This multi-step transformation is essential for parasite survival and pathogenesis, conferring *P. falciparum* its ability to maintain chronic infections in humans. A large fraction of exported proteins are recognizable by the presence of a 5-amino acid motif, known as the *Plasmodium* export element or PEXEL^10,11^, while others have no discernable primary sequence motif and are termed as PEXEL-negative exported proteins or PNEPs^12^. Most PNEPs possess a transmembrane (TM) domain that serves to target them to the ER and the secretory pathway^12^. Several of these PNEPs play a critical role in malaria pathogenesis, such as skeleton-binding protein 1 (SBP1)^13–15^, membrane associated histidine-rich protein (MAHRP1)^16,17^ and erythrocyte membrane protein 1 (PfEMP1)^18–20^.

Exported membrane proteins are inserted into ER membrane during their synthesis^12,21,22^. These membrane proteins are transported via vesicles from the ER and inserted into the parasite plasma membrane (PPM) when the transport vesicles fuse to the PPM^22^. While it has been shown that all exported proteins require the *Plasmodium* translocon of exported proteins (PTEX) to cross the PV membrane (PVM)^23,24^, how membrane proteins are extracted from the PPM and delivered to the PTEX complex remains unknown. It has been postulated that a putative *Plasmodium* translocon of exported membrane proteins (which we term as PTEM)^25,26^ is required for extraction of membrane proteins from the PPM either alone or in cooperation with the PTEX unfoldase HSP101^27,28^ (Fig. 1A). The identity of proteins in this putative PTEM complex are unknown and there are no obvious candidates in the genome of *P. falciparum*. Therefore, we attempted to develop an unbiased proteomic approach to identify proteins that could form a putative PTEM complex.

**Figure 1.**
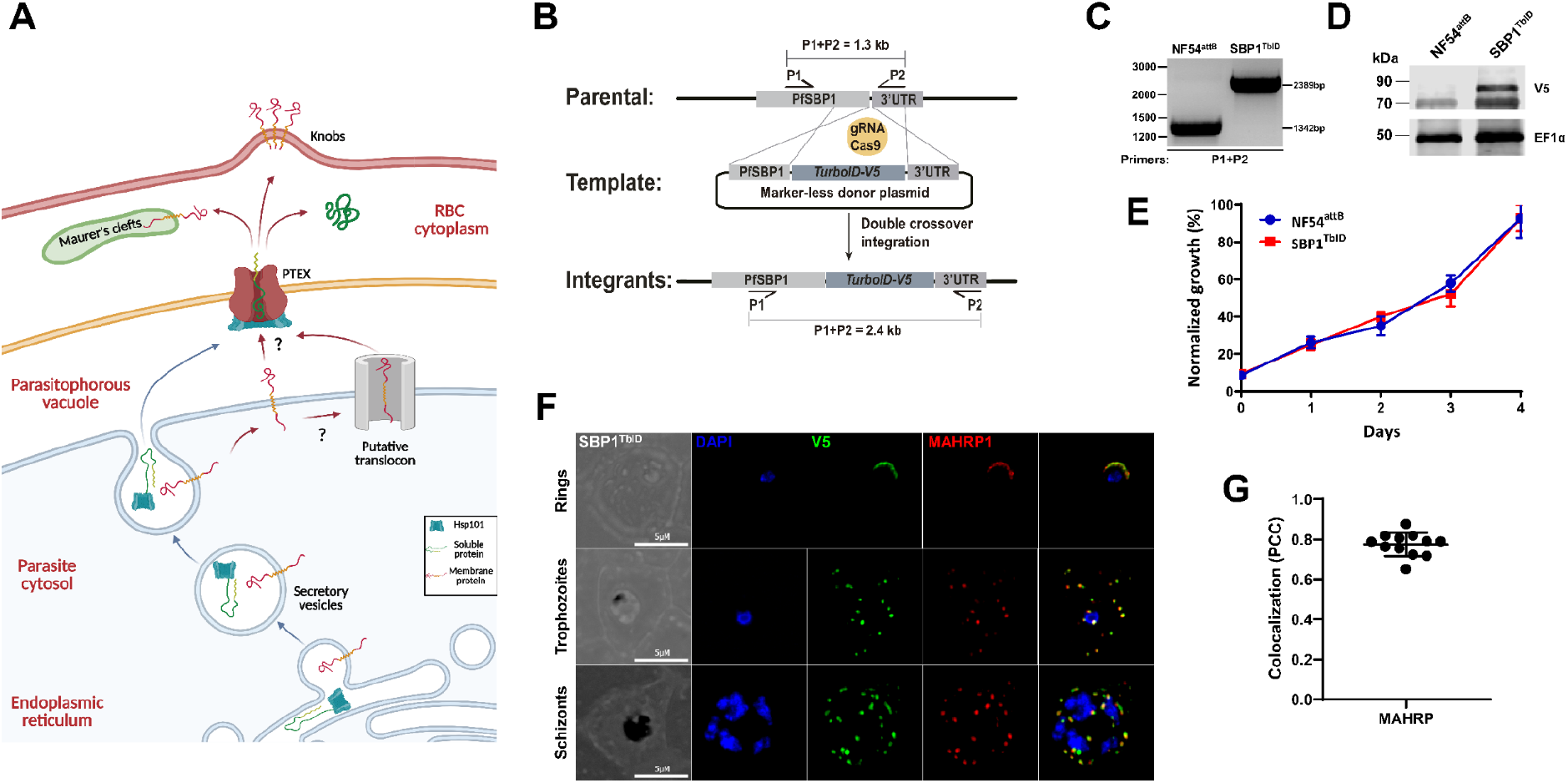
Generation of SBP1^TbID^ mutants. (A) Schematic of protein export. Membrane and soluble proteins are transported from the ER into the PV by secretory vesicles. Soluble proteins are released into the PV lumen after fusion of the secretory vesicle to the PPM. Membrane proteins, on the other hand, are inserted into the PPM and need to be extracted from the membrane by a putative translocon (PTEM) for further transport through the PV membrane to the RBC cytoplasm by the PTEX complex. Soluble and membrane proteins are transported to their final location in the infected RBC. (B) Schematic showing the integration of the repair plasmid used to tag the genomic loci of SBP1 with TurboID-V5. Cas9 introduces a double-stranded break at the C-terminus of the SBP1 locus. The repair plasmid provides homology regions for double-crossover homologous recombination, introducing TurboID and the V5 tag sequences. (C) PCR test confirming integration at the SBP1 locus. Amplicons were amplified from genomic DNA isolated from mutant and wild-type parasites. Primers were designed to amplify the region between the C-terminus and the 3’UTR of SBP1. All primers are in Supplemental Table 1. (D) Western blot of parasite lysates isolated from the parental line (NF54^attB^)^36^ and a clone of SBP1^TbID^ (D10) probed with antibodies against V5 and EF1α (loading control). The protein marker sizes are shown on the left. (E) Growth of asynchronous SBP1^TbID^ parasites, compared to the parental line NF54^attB^, over 4 days via flow cytometry. 100% represent the highest value of calculated parasitemia. Representative of three biological replicates shown for each growth curve. Each data point represents the mean of three technical replicates; error bars represent standard deviation. (F) IFA showing SBP1^TbID^ localizes to the parasite periphery in early-ring stage (top) and is exported to the Maurer’s cleft in trophozoite (middle) and schizont (bottom) stages. Asynchronous SBP1^TbID^ parasites were fixed with acetone and stained with specific antibodies. Images from left to right are phase-contrast, DAPI (nucleus, blue), anti-V5 (green), anti-MAHRP (red), and fluorescence merge. Z stack images were deconvolved and projected as a combined single image. (G) Quantification of the colocalization of SBP1^TbID^ with MAHRP using the Pearson’s correlation coefficient. Three biological replicates represented with 4 late-stage-parasite images from each replicate. Error bars represent standard deviation.

Proteomic approaches have been used previously to identify the exported-protein interacting complex (EPIC) at the PV, which is thought to be required for protein export^29^. Similar approaches using *Plasmodium* exported proteins have identified stable complexes at the Maurer’s clefts (MC), a parasite-generated protein sorting organelle in the RBC^30,31,32,33^. However, the identification of the putative PTEM has proven elusive because its interaction with exported membrane proteins will be transient and therefore, unlikely to be captured using immunoprecipitation assays which are heavily biased towards stable complexes. Therefore, we used a rapid, proximity-labeling approach to attempt to identify a putative PTEM complex. To our knowledge, this approach has not yet been used in a time-resolved manner to capture transient interactions in the secretory pathway.

We chose to tag the endogenous SBP1 gene (PF3D7_0501300) with a new iteration of the promiscuous biotin ligase BirA, known as TurboID (generating SBP1^TbID^)^34^. SBP1 is a PNEP with a single transmembrane domain and is exported in early ring-stage parasites to the MC^13^. TurboID is a highly efficient enzyme that biotinylates proteins in proximity within 10 minutes^34^. Therefore, we hypothesized that SBP1^TbID^ will biotinylate proteins, even those transiently interacting with SBP1^TbID^ along the secretory pathway during its export to the MC. Since SBP1^TbID^ should rapidly biotinylate proximal proteins, we further reasoned that we could differentiate early (pre-export) interactors from late (post-export) interactors of SBP1^TbID^. Our data show that the SBP1^TbID^ fusion protein is exported to the MC efficiently and with similar kinetics to another MC protein, MAHRP1. Critically, SBP1^TbID^ can rapidly biotinylate proximal proteins prior to its export from the PV as well as after export at the MC. Using label-free quantitative proteomics, we compared pre-export interactors and post-export interactors of SBP1^TbID^. This approach led to the identification of two membrane associated proteins that may be part of the putative PTEM complex.

## RESULTS

### SBP1 fused to TurboID is exported to Maurer’s Clefts

Using CRISPR/Cas9 gene editing we generated mutants of SBP1 (Fig. 1B), where the endogenous gene was tagged with the TurboID biotin ligase (SBP1^TbID^)^34,35^. We chose TurboID because it is an optimized version of the biotin ligase BirA^34^. TurboID is a highly active mutant of BirA with an increased biotinylation radius and faster biotinylation kinetics^34,35^. PCR analysis of genomic DNA isolated from the SBP1^TbID^ parasite line showed the correct integration of the TurboID biotin ligase and a V5 tag at the endogenous locus of SBP1 (Fig. 1C). We detected expression of the SBP1^TbID^ in the mutant line at the expected size, but not in the parental line (Fig. 1D). To ensure that the expression of TurboID is not detrimental to the parasite, we observed the growth of SBP1^TbID^ and the parental parasite line (NF54^attB^)^36^, over several asexual cycles using flow cytometry (Fig. 1E). These data show no difference in the asexual growth of SBP1^TbID^ compared to the parental parasites, demonstrating that expression of TurboID or its fusion to SBP1 does not inhibit parasite growth.

SBP1 is an exported protein with a single transmembrane domain synthesized in the parasite ER and transported to the MC in the RBC cytoplasm^37,38^. To confirm that fusing TurboID to SBP1 did not inhibit its export to the MC, we used immunofluorescence microscopy (IFA) to test whether SBP1^TbID^ colocalized with another MC resident protein, MAHRP1^17^. Our data show that SBP1^TbID^ is exported from the parasite to the MC and co-localizes with MAHRP1 during trophozoite and schizont stage parasites (Fig. 1F and 1G). On the other hand, in early ring stage parasites, we show that SBP1^TbID^, as well as MAHRP1, localize to the periphery of the parasite, probably in the PV prior to export (Fig. 1F). SBP1 accumulation in electron dense regions within the parasite plasma membrane (PPM) before transport through the PV membrane has previously been observed by electron microscopy ^39^. These data suggest that SBP1 and possibly other MC resident proteins accumulate in the PV before being exported to the infected RBC. Together, our data show that tagging SBP1 with the TbID biotin ligase did not alter the asexual growth or development of the parasite, nor did it inhibit the export of SBP1 to the host RBC and MC (Fig. 1).

### Biotin-dependent proximity labeling by SBP1^TbID^

Since our data show that the SBP1^TbID^ fusion protein was exported to MC, we wanted to examine the capacity of TurboID to biotinylate proximal proteins in SBP1^TbID^ parasites. TurboID is an extremely efficient enzyme and we observed it could utilize the minimal amount of biotin present in the media used to grow SBP1^TbID^ parasites (Fig. S1). The ability of SBP1^TbID^ to use the biotin present in complete media meant that biotinylation of proximal proteins would occur constantly, which would prevent the differentiation of early interactors from late interactors. Thus, we grew SBP1^TbID^ in minimal media without biotin because the normal asexual development of *P. falciparum* does not require biotin^40^. To test if SBP1^TbID^ biotinylation is dependent upon the presence of exogenous biotin, we analyzed protein extracts of asynchronous parasites in the presence or absence of biotin by streptavidin blotting. We observed that efficient biotinylation of proximal proteins occurs only in the presence of biotin in SBP1^TbID^ parasites (Fig. 2A). Self-biotinylation in SBP1^TbID^ parasites was observed in the presence or absence of biotin (Fig. 2A, lane 3 and 4, see asterisk), in agreement with previously reported when tagging proteins with TurboID^34,41^. No endogenous biotinylation was detected in the parental line NF54^attB^, showing that biotinylation occurs only when TurboID is being expressed by the parasite line (Fig. 2A, lane 1 and 2). These data show that SBP1^TbID^ efficiently biotinylates proteins and its activity is dependent upon the presence of biotin in the growth medium.

**Figure 2.**
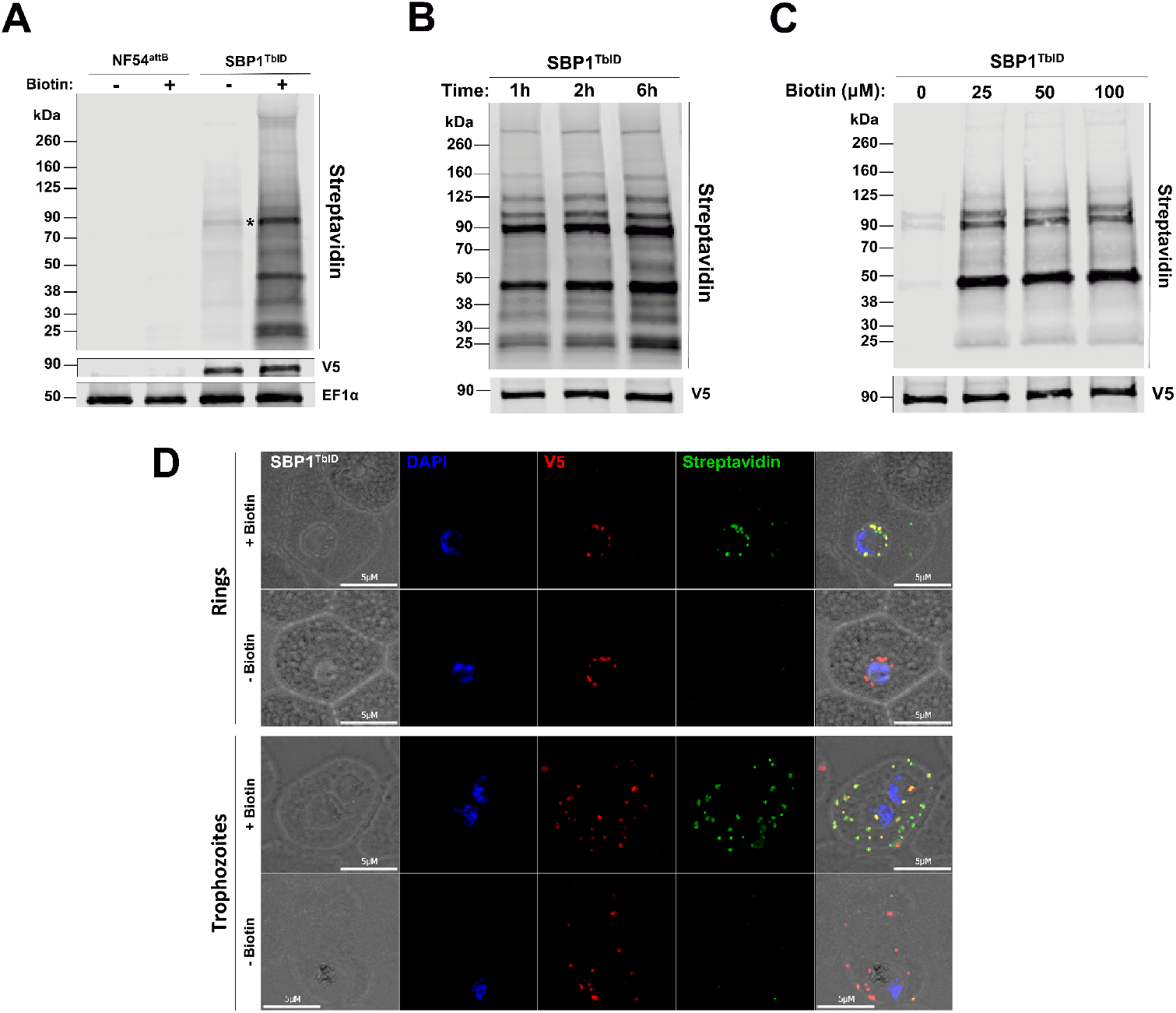
Biotinylation of proximal proteins by TurboID_V5_-tagged SBP1. (A) Western blot of parasite lysates isolated from the parental line NF54^attB^ and the mutant line SBP1^TbID^ incubated with or without biotin (50 μM) for 2 h. Samples were probed with antibodies against V5, EF1α (loading control) and fluorescent dye-labeled streptavidin. The protein marker sizes are shown on the left. (B) Western blot of parasite lysates isolated from the mutant line SBP1^TbID^ incubated with biotin (50 μM) for 1 h, 2h and 6h. Samples were probed with antibodies against V5 (loading control) and fluorescent dye-labeled streptavidin. The protein marker sizes are shown on the left. (C) Western blot of parasite lysates isolated from the mutant line SBP1^TbID^ incubated with different concentrations of biotin (0, 25, 50 and 100 μM) for 1 h. Samples were probed with antibodies against V5 (loading control) and fluorescent dye-labeled streptavidin. The protein marker sizes are shown on the left. (D) IFA showing SBP1^TbID^ biotinylates proteins during their export out of the parasite (top panels) and at their final location at the Maurer’s clefts (Bottom panels). Asynchronous SBP1^TbID^ parasites were fixed with acetone after 2 h of incubation with biotin (50 μM) and stained with specific antibodies. Images from left to right are phase-contrast, DAPI (nucleus, blue), anti-V5 (red), streptavidin (green), and fluorescence merge. Z stack images were deconvolved and projected as a combined single image.

TurboID is a highly active enzyme^34^ that offers the possibility of rapid and time-resolved labeling approaches in contrast to previous proximity-labeling methods with much longer incubation times, usually greater than 12 hours^42–44^. Thus, we wanted to assess whether SBP1^TbID^ is able to rapidly biotinylate proximal proteins. SBP1^TbID^ parasites were incubated with biotin for 1, 2 or 6 hours and the biotinylation of proteins were observed using western blots probed with streptavidin (Fig. 2B). We also tested biotinylation in response to different concentrations of biotin, 25, 50 and 100 μM (Fig. 2C). Biotinylated proteins were observed at all time points and biotin concentration, and the observable difference in the extent of protein biotinylation between the time points and concentrations was minimal (Fig. 2B and 2C).

The SBP1^TbID^ fusion protein has to traverse several membranes during its export to the MC, and therefore, it is likely to unfold and then refold during this transport process. Furthermore, to our knowledge TurboID has not yet been utilized in a time-resolved manner to identify transient interactors as proteins are transported through the secretory pathway. Therefore, we wanted to determine if SBP1^TbID^ parasites could biotinylate proteins proximal to SBP1 at different cellular locations during the export of SBP1^TbID^ from the parasite ER to the MC. Synchronized early ring and trophozoite stage parasites were observed by IFAs after the addition of biotin for 2 h. We observed biotinylation at the parasite periphery, possibly when SBP1^TbID^ accumulates at the PV^39^ (Fig. 2D, top panels). Biotinylation was also observed when SBP1^TbID^ had been exported to the MC (Fig. 2D, bottom panels). The observed biotinylation was dependent upon the addition of biotin. Together, these data demonstrate that SBP1^TbID^ was highly active, efficient, rapid, and labeled proximal proteins at different subcellular locations during its export from the parasite ER to the final location at the MC.

### Early interactors of SBP1^TbID^ identified by proximity labeling

Since our data show that SBP1^TbID^ biotinylates proximal proteins during its transport from the parasite ER into the RBC cytoplasm (Fig. 2C), we next wanted to identify the *P. falciparum* effectors that interact with SBP1 at the host-parasite interface. To do so, we wanted to define the kinetics of SBP1^TbID^ transport from its site of synthesis in the parasite ER to the MC and test if we could reproducibly detect SBP1 at the host-parasite interface. As previously described (Fig. 1E, 2D), SBP1^TbID^ and proteins biotinylated by SBP1^TbID^ could be detected at the parasite-RBC interface. To assess whether we could reproducibly observe SBP1^TbID^ within the parasite prior to its export to the host RBC, we observed the subcellular localization of SBP1^TbID^ with respect to EXP2, a PVM resident protein^46,47^, at different time points after parasite invasion in tightly synchronized cultures. SBP1 has been detected at the MCs as early as 4-6 hours post invasion (hpi)^48^, therefore, we observed the subcellular location of SBP1^TbID^ in parasites at 3, 4 and 5 hpi. In some SBP1^TbID^ parasites, SBP1 was either not detectable or not expressed (Fig. 3B, top panels). As expected, we found parasites where SBP1^TbID^ was within the PV periphery, and others where the protein was already exported to the RBC cytoplasm (Fig. 3C, mid and bottom panels). We quantified these three events over several biological replicates. At 4hpi, SBP1^TbID^ was not expressed in about 30% of the parasites, exported in ∼10% of observed parasites and at the host-parasite interface in the vast majority (60%) of all parasites (Fig. 3C). These data showed us that harvesting proteins biotinylated by SBP1^TbID^ at the host-parasite interface would be feasible.

**Figure 3.**
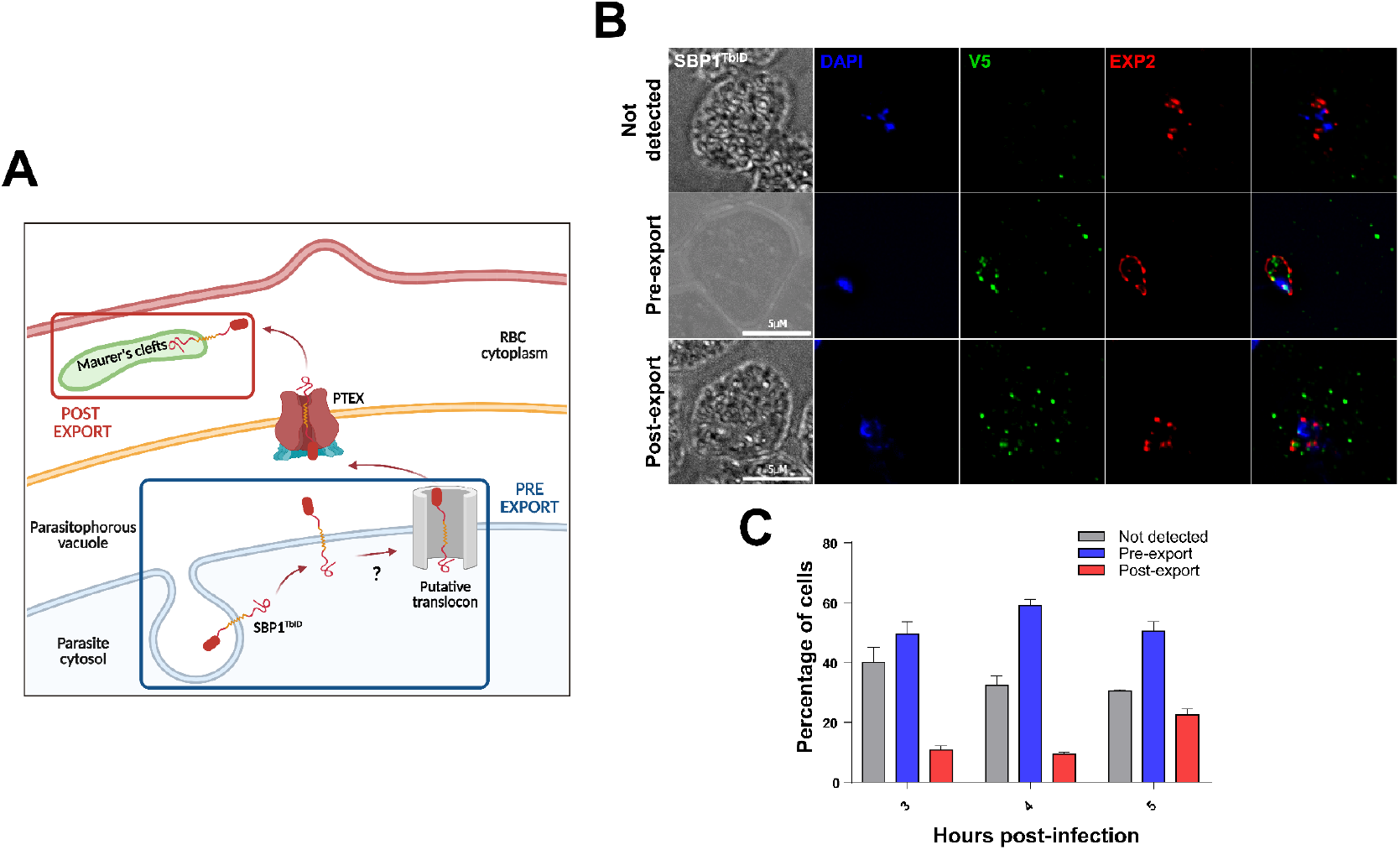
(A) Schematic of the export of proteins in *P. falciparum* highligting the locations where proteins biotinylated by SBP1^TbID^ will be harvested. Created with BioRender.com. (B) IFA showing the different localizations of SBP1^TbID^ during its export at early ring-stages (3-5 hpi). Tightly synchronous SBP1^TbID^ parasites were fixed with acetone at 3 h, 4h and 5 h post infection, and stained with specific antibodies. Images from left to right are phase-contrast, DAPI (nucleus, blue), anti-V5 (green), EXP2 (PV marker, green), and fluorescence merge. Z stack images were deconvolved and projected as a combined single image. (C) Quantification of the events observed in (B). Events were scored based on the localization of SBP1^TbID^ with respect to the PV marker EXP2. A total of 50 parasites were scored for each time point. n= 3 biological replicates; error bars represent standard deviation.

To identify early interactors of SBP1, especially those at the host-parasite interface, we opted for a quantitative and comparative approach. We wanted to differentiate these early interactors from SBP1 interactors at the MC, which have been previously identified^32^, as well as those being co-transported with SBP1 to the MC. We hypothesized that using label-free quantitative proteomics and comparing interactors isolated from 4 hpi and 20 hpi would allow us to identify the early interactors of SBP1. By 20 hpi, all SBP1 is at the MCs and no more SBP1 is synthesized^49^. Label-free proteomics have been shown to offer a large dynamic range and high proteome coverage in the identification of biotinylated proteins^41,50–52^.

First, tightly synchronized late-stage schizonts were collected. These parasites were then split into two samples, one incubated with biotin for 4 h until 4 hpi (Fig. 3A, blue square) and then collected for further processing. Since our data show that in the majority of the 4 h ring stage parasites, SBP1^TbID^ was at the host-parasite interface (Fig. 3C), parasites were incubated with biotin for 4 h to maximize the labeling of proximal proteins and capture a larger fraction of the pre-export interactors. To collect the post-export sample, we let SBP1^TbID^ parasites develop until 16hpi because all SBP1 localizes to MC by this time and this protein is no longer synthesized. Thus, this sample was allowed to develop without biotin for 16h, and then incubated with biotin for 4 h until 20 hpi (Fig. 3A, red square). Biotinylated proteins were isolated from parasite lysates using streptavidin-affinity pulldown. Streptavidin-captured proteins were identified via mass spectrometry (MS) and quantified over several biological replicates^41,50,51^ (Fig. 4A). A total of 1,122 proteins were identified in at least one of the replicates. We then compared the proteins identified in the 4h sample with those identified in the 20h sample (Fig. 4B). We defined the putative pre-export interactors of SBP1 from our dataset using three stringent criteria. Proteins enriched more than 10-fold compared to the 20 hpi samples with a p-value cut-off of 0.05, and present in all three biological replicates were considered as differentially labeled interactors at 4 hpi. Using these criteria, 24 protein candidates were identified as putative pre-export interactors of SBP1^TbID^ during its transport at the parasite-RBC interface (Fig. 4B). The identified proteins were classified into subgroups based on their predicted functions and subcellular locations^53^. Of the 24 identified proteins, 11 were uncharacterized proteins with no predicted function. As expected, this approach identified proteins known to be involved in protein and vesicle transport (5/24). One of the statistically significant interactors of SBP1 was EXP3 (3-fold enriched at 4hpi), which has been localized to the PV and functions in protein export ^29^. SBP1 (star, Fig. 4B) and other MC localized proteins were also identified but were not enriched at either time point, demonstrating that the experiment worked as designed (Fig. 4B). Identification of other exported proteins, including some MC proteins, only in the post-export (20hpi) samples further suggests that the proteomic approach using SBP1^TbID^ worked as designed. Together these data show that our approach successfully identified a group of proteins differentially biotinylated by SBP1^TbID^ prior to its export to the MC.

**Figure 4.**
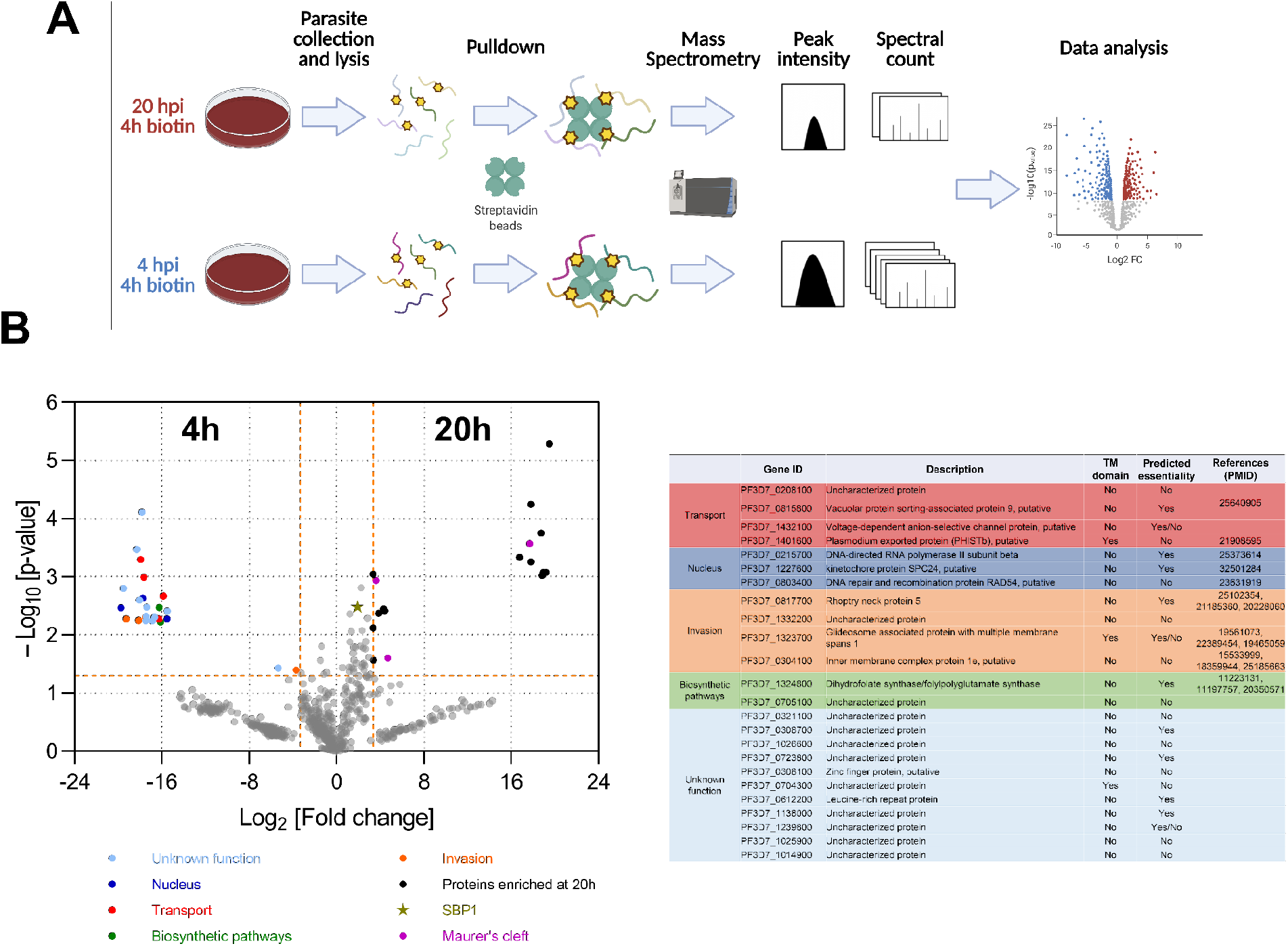
(A) Schematic of the experimental design for time-resolved biotinylation and proteomics to identify pre-export and post-export interactors of SBP1^TbID^. Created with BioRender.com. (B) Interactors enriched at 4 hpi (p-value plotted as function of fold change between the two samples). Proteins with p-value ≤0.05 and more than 10-fold change are identified as SBP1^TblD^ interactors. n= 3 biological replicates. (C) A summary table of the putative interactors of SBP1^TbID^ at 4hpi grouped by their putative functions are shown. All proteins identified are in Supplemental Table 2.

### Early interactors of SBP1^TbID^ localize to the host-parasite interface

Since we were interested in identifying proteins that may act as a putative translocon to extract exported membrane proteins at the PPM, we reasoned that membrane associated proteins among pre-export SBP1 interactors could function in this role. Thus, based on membrane-association, high statistical score and fold enrichment, we selected the Glideosome-associated protein with multiple membrane spans 1 (GAPM1, PF3D7_1323700) as one putative candidate. GAPM1 is a membrane protein associated with the biogenesis of the Inner Membrane Complex (IMC) in asexual and sexual stages. GAPM1, as part of the IMC, is suggested to have a role in merozoite invasion^54–56^. Using these criteria, another putative candidate was the channel protein Voltage-dependent anion-selective channel protein (VAC, PF3D7_1432100). VAC is a soluble protein with a translocon of outer mitochondrial membrane (TOM40) domain but no mitochondria targeting signal. Nothing is known about the function of VAC in *P. falciparum*.

To characterize these proteins, we used CRISPR/Cas9 gene editing to generate the conditional mutants, termed VAC^apt^ and GAPM1^mNG-apt^. In these parasite lines, their endogenous loci were tagged with the *tetR* aptamer system, which results in anhydrotetracycline (aTc)-dependent expression of protein (Fig. 5A and 5B)^57^. PCR analysis of genomic DNA from VAC^apt^ and GAPM1^mNG-apt^ parasite lines showed correct integration of the knockdown system at the endogenous loci (Fig. S2A-C). To assess the efficiency of the knockdown system, we measured protein expression in the presence or absence of aTc by Western blotting. For both proteins, there is a clear reduction of protein expression (Fig. S2D). Knockdown of GAPM1 was detrimental, as parasites were not able to progress into a second life cycle (Fig. S2E). Knockdown of VAC also inhibited the asexual expansion of VAC^apt^ parasites (Fig. S2E).

**Figure 5.**
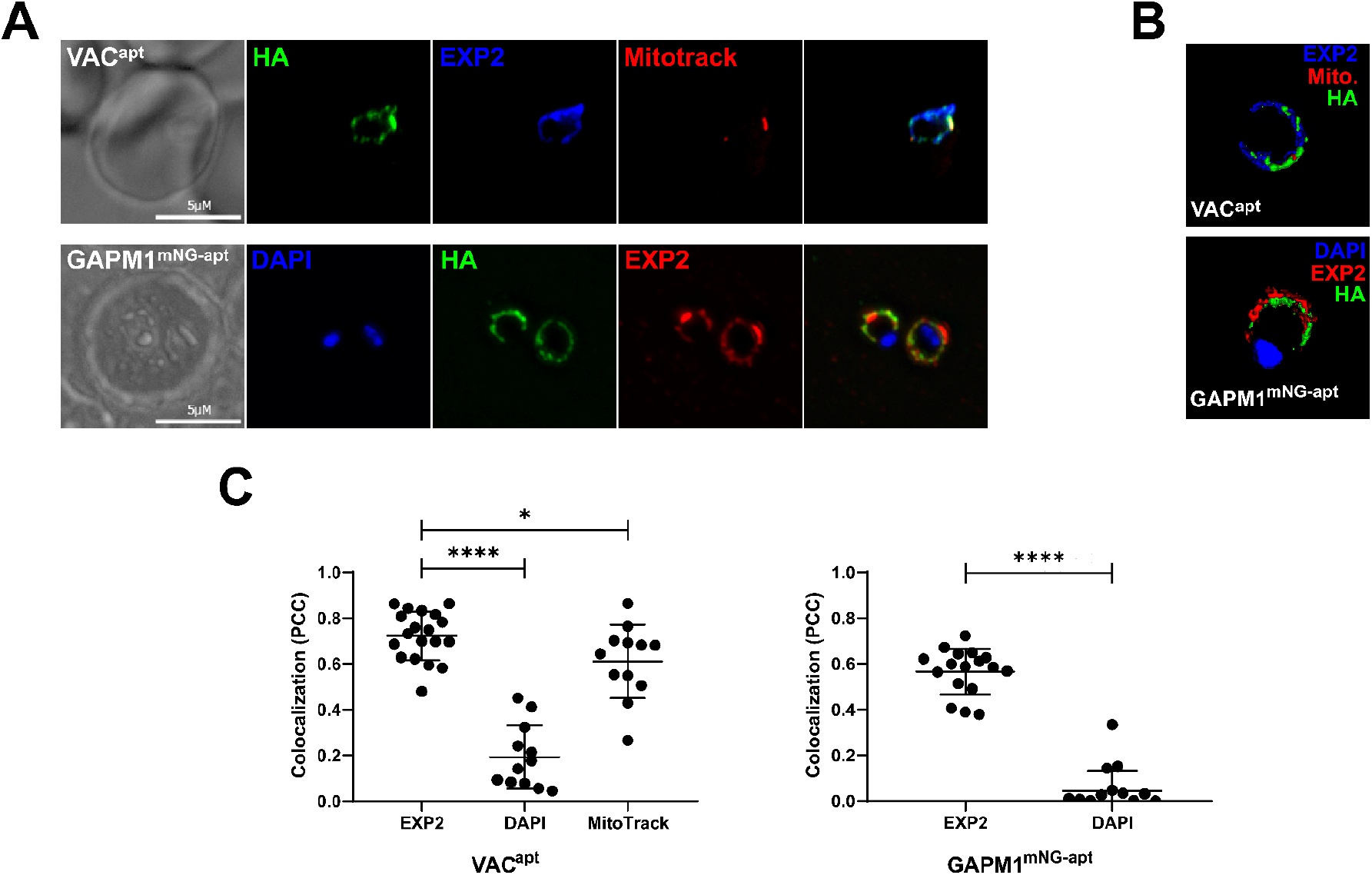
Generation and characterization of VAC^apt^ and GAPM1^mNG-apt^. (A) Representative IFA showing VAC^apt^ and GAPM1^mNG-apt^ localizes within the PV in the early-ring stage (4 hpi). Tightly synchronous parasites were fixed with PFA (VAC^apt^) and acetone (GAPM1^mNG-apt^) and stained with specific antibodies. Images of VAC^apt^ from left to right are phase-contrast, anti-HA (green), anti-EXP2 (PV, blue), mitotracker (mitochondria, red), and fluorescence merge. Images of GAPM1^mNG-apt^ from left to right are phase-contrast, DAPI (nucleus, blue), anti-HA (green), anti-EXP2 (PV, red), and fluorescence merge. Z stack images were deconvolved and projected as a combined single image. (B) 3D reconstruction based on structured illumination microscopy images captured from VAC^apt^ and GAPM1^mNG-apt^ ring-stage parasites at 4 hpi and stained with the antibodies as in (A). (C) Quantification of the colocalization of VAC and GAPM1^mNG^ with respect to EXP2, MitoTracker (Mitochondrial marker) and DAPI, and to EXP2 and DAPI, respectively, using the Pearson’s correlation coefficient. Three biological replicates represented with 6 parasite images from each replicate for EXP2, and 3 parasite images for MitoTrack and DAPI. Error bars represent standard deviation. ****p<0.05 by t-student test.

Our data show that GAPM1^mNG^ localizes to the IMC in schizonts (Fig. S3). However, what happens to the IMC after merozoite invasion is unclear and we reasoned that the IMC most likely fuses to the PPM shortly after invasion. Similarly, the subcellular localization of VAC during the early stages of the asexual lifecycle was unknown. The proteomic data suggest that these proteins are proximal to SBP1^TbID^ when SBP1 is in the PV (Fig. 4B). Therefore, we used IFAs to localize both proteins in tightly synchronized parasites at 4 hpi with respect to the PV marker EXP2. VAC^apt^ localizes to the parasite periphery and is closely juxtaposed with the known PV marker, EXP2, but it also partially overlaps with the mitochondria (Fig. 5A, top panels) suggesting that it may be localized to both subcellular organelles. GAPM1^mNG-apt^ localizes to the parasite periphery at 4 hpi, and shows colocalization with EXP2 (Fig. 5A, bottom panels). To corroborate our observations by IFAs, we used structured illumination microscopy (SIM) to determine the subcellular localization of VAC and GAPM1. Both proteins are closely juxtaposed to the PVM marker EXP2, which suggests VAC and GAPM1 do not localize to the PVM (Fig. 5B). The SIM data show that both VAC and GAPM1 follow the same contours as EXP2 but the signals do not completely overlap, which suggests that VAC and GAPM1 localize to another membrane (Fig. 5B and Fig. S4). Together these results show that both GAPM1 and VAC localize to the same region together with SBP1^TbID^ at the PV at 4 hpi, as suggested by the proximity labeling data.

## DISCUSSION

The protein-protein interactions that usher exported proteins to their final destinations in the RBC via the secretory pathway are transient in nature. Previously, IP-based methods have been used to identify proteins required for export of *P. falciparum* proteins, such as the PTEX complex^58^ and the EPIC complex^29^. While IP-based approaches are well-suited for identifying stable complexes, they are unlikely to identify transient interactions. Consequently, a putative additional translocon at the PPM required for extracting exported membrane proteins which are inserted into the PPM during transport has long been proposed^25,26^, but no candidates for this putative PTEM complex have been identified as yet (Fig. 1).

To identify candidates for a putative PTEM complex, we used time-resolved biotinylation to identify transient interactions of an exported membrane protein, SBP1, during its export. This approach uses a rapid and promiscuous biotin ligase to biotinylate proximal proteins^34^. Since biotinylation is a permanent modification, even transient interactions can be potentially identified. Our data show that fusion of TurboID to the exported transmembrane-containing protein, SBP1, did not alter its trafficking to the MC nor did it have any effect on parasite growth. These data also suggest that TurboID is enzymatically active during transit in the parasite secretory pathway.

A previous study on the SBP1 interactome at their final location at the Maurer’s cleft identified 88 parasite proteins as putative interactors^32^. Most of the top-ranked hits from the previous study were also identified in our study, including PfEMP1, Pf332, PIESP2, REX1, MAHRP1, PTP1 and vapA, but were not highly enriched (≤10-fold) in the post-export interactors fraction. It is possible that some of these proteins are co-transported with SBP1 and thus, are identified in the pre-export fraction as well. Members of the PTEX complex such as EXP2, HSP101, and Trx2, were also identified in the pre-export fraction, albeit below statistical significance. In addition, PTP2 and PfG174, which have previously been shown to localize as residents^59^, or transient interactors^60^ of the Maurer’s clefts, were more than 10-fold enriched at the post-export time point, demonstrating the reliability of our approach to identify SBP1 interactors. Another subset of proteins identified in our study as post export interactors of SBP1 are ribosomal proteins, which have been previously observed to be exported to the red blood cell cytoplasm in *P. falciparum*^*61*^. Together, these data strongly suggest that the time-resolved, rapid biotinylation approach was working as designed. Since our focus was to identify pre-export interactors, we did not pursue these proteins for further study.

Using label-free quantitative proteomics, we identified a group of 24 putative candidates that interact with SBP1 prior to its export to the RBC. Several of the proteins identified (11/24) were uncharacterized proteins. Since we undertook this approach to identify the proposed translocon of exported membrane proteins, we did not pursue the function of these proteins in this study. Translocons function to transport proteins across membranes and therefore, we hypothesized that membrane associated proteins in this list could putatively function as translocons. There were two putative candidates in the pre-export interactors of SBP1 that were membrane associated, VAC and GAPM1. However, their localization in early ring stage parasites was unknown. Therefore, to study the function of VAC and GAPM1 in early ring stage parasites, we successfully generated conditional mutants. The data show that both VAC and GAPM1 play important functions in parasite survival within the infected RBC. Knockdown of these proteins inhibits parasite growth. However, achieving protein knockdown takes about 48 hours and results in parasite death prior to re-invasion of the RBC. Therefore, this prevents the characterization of their role in export, which occurs in about 6-8 hours after invasion. Similar to the PTEX translocon, EXP2^46^, it is likely that both GAPM1 and VAC have other essential functions in the asexual lifecycle. Defining their function in export will require using a more rapid knockdown approach with similar kinetics as SBP1 export, such as degradation-domain based tools^23,62^ or rapid mislocalization based methods^63^.

VAC has a β-barrel porin domain that can form an aqueous channel in the membrane and function as a translocon in mitochondria and other plastids^64^. In a recent proximity-biotinylation based proteomic screen to catalog mitochondrial proteins in *P. falciparum*, VAC was pulled down in the membrane fraction of parasite lysates, and not in the mitochondrial fraction^65^. In addition, VAC does not have a predicted mitochondrial targeting sequence, in contrast to its ortholog, TOM40 (PF3D7_0617000), which has a mitochondrial targeting sequence, suggesting that VAC might not be localized to the mitochondria^66^. Our data reveal that VAC localizes at the host-parasite interface in early ring stages. While there is some overlap of VAC with the mitochondria, there is stronger overlap between the PVM marker, EXP2, and VAC in lower resolution IFAs. It is also possible that VAC is dually localized both to the mitochondria as well as to the host-parasite interface. SIM data suggest that EXP2 and VAC are closely juxtaposed but with minimal overlap. This suggests that VAC localizes to a compartment in close proximity to the PVM, most likely the PPM. On the other hand, GAPM1 has seven TM domains and is from an apicomplexan-specific family of proteins^56^. GAPM1 has been localized to the inner membrane complex (IMC) in schizont-stage parasites^56^. The IMC plays an essential role in the invasion of merozoites into the RBC, however, it is unclear what happens to the IMC post-invasion. Lower resolution IFAs show that GAPM1 co-localizes with the PVM localized EXP2 in early ring stage parasites. Using SIM, we observed GAPM1 in close juxtaposition with EXP2, but they do not completely overlap, suggesting that GAPM1, like VAC, localizes to a membrane compartment at the parasite periphery that is not the PVM. Given these data, it is likely that GAPM1 may also localize to the PPM in early rings. These data further suggest that the IMC may fuse to the parasite plasma membrane after merozoite invasion. Together, these findings suggest that VAC and GAPM1 transiently interact with SBP1 prior to its export and their localization to the parasite periphery suggests a role in export, perhaps as members of a putative PTEM complex. Based on these data, we propose a model where GAPM1 and VAC act together to select and extract exported membrane proteins from the PPM for further transport via the PTEX into the host RBC.

Several mechanistic aspects of this model remain to be resolved but like the PTEX complex, which was first identified as a putative complex at the PV membrane^58^. Both VAC and GAPM1 were reproducibly biotinylated by SBP1^TbID^ prior to its export and localize to the parasite periphery. Further, VAC has a porin translocon domain which could function in a manner analogous to the mitochondrial outer membrane translocon to extract membrane anchored exported proteins from the parasite plasma membrane. This hints that this porin domain protein has been repurposed by *Plasmodium* parasites on the PPM to facilitate export of membrane proteins to the infected RBC.

## ACKNOWLEDGEMENTS

We thank Chi-Min Ho for helpful comments on the manuscript; Dan Goldberg for anti-EF1α; Hans-Peter Beck for anti-MAHRP1; the European Malaria Reagent Repository for anti-EXP2; Julie Nelson at the CTEGD Cytometry Shared Resource Laboratory for help with flow cytometry and analysis; and Muthugapatti Kandasamy at the Biomedical Microscopy Core at the University of Georgia for help with microscopy. We acknowledge the assistance of Phil Gafken at the Proteomics Resource at Fred Hutchinson Cancer Research Center for mass spectrometry analysis. The study was funded by NIH/NIAID R01AI130139 (V.M.), T32AI060546 (D.W.C.), and CTEGD

Fellowship funded by Provost’s Office at UGA (D.A.).

## MATERIALS AND METHODS

### Construction of SBP1 plasmids

Genomic DNA was isolated from *P. falciparum* NF54^attB^ cultures using the QIAamp DNA blood kit (Qiagen). PCR products were inserted into the respective plasmids using ligation-independent cloning (SLIC), as described earlier ^67^, or the NEBuilder HiFi DNA Assembly system (NEB). All constructs used in this study were confirmed by sequencing. All primers used in this study are in Supplemental Table 1.

For generation of the plasmid pTOPO-SBP1-TbID, sequences of approximately 500 bp of homology to the SBP1 C-terminus and 3’UTR were amplified using primer pairs P1-P2 and P3-P4, respectively, and the sequence of V5-tagged TurboID was amplified using primers P5 and P6. For expression of a SBP1 gRNA, oligos P17-P18 were inserted into cut pUF1-Cas9.

For generation of the plasmid pKD-VAC-Apt, sequences of approximately 450 bp of homology to the Pf1432100 C-terminus and 3’UTR were amplified using primer pairs P7-P8 and P9-P10, respectively. Amplicons were then inserted into pKD^57,67^ digested with AatII and AscI. For expression of a Pf1432100 gRNA, oligo P19 was inserted into cut PUF1-Cas9.

For generation of the plasmid pKD-GAPM1-mNG-Apt, sequences of approximately 500 bp of homology to the PfGAPM1 C-terminus and 3’UTR were amplified using primer pairs P11-P12 and P13-P14, respectively, and the sequence of mNeonGreen was amplified using primers P15 and P16. Amplicons were then inserted into pKD^57^ digested with AatII and AscI. For expression of PfGAPM1 gRNA, oligo p20 was inserted into cut PUF1-Cas9.

## Parasite culture and transfections

*Plasmodium* parasites were cultured in RPMI 1640 medium (NF54^attB^, VAC^apt^ and GAPM1^mNG-apt^) or in biotin-free medium (SBP1^TbID^, VAC^apt^/SBP1^TbID^) ^68^ supplemented with AlbuMAX I (Gibco), and transfected as described earlier^69^.

For generation of SBP1^TbID^ parasites, a mix of two plasmids (50 µg each) were transfected into NF54^attB^ parasites in duplicate. The plasmid mix contained the plasmid pUF1-Cas9-SBP1gRNA, which contains the DHOD resistance gene, and the marker-free plasmid pTOPO-SBP1-TbID. Drug pressure was applied 48 h after transfection, using 1µM DSM1 ^70^ and selecting for Cas9 expression. After parasites grew back from transfection, integration was confirmed by PCR, and then cloned using limiting dilution. After clonal selection, cultures were transferred to biotin-free medium without DSM1.

For generation of VAC^apt^ and GAPM1^mNG-apt^ parasites, the pKD-VAC-Apt and pKD-GAPM1-mNG-Apt plasmids (20 µg) and the respective pUF1-Cas9 plasmid (50 µg) were transfected into NF54^attB^ parasites in duplicate. Before transfection pKD plasmids were digested overnight with EcoRV (NEB). The enzyme was then subjected to heat inactivation for 20 min at 65 °C and then mixed with the pUF1-Cas9 plasmid.

Transfected parasites were grown in 0.5 µM anhydrous tetracycline (aTc) (Cayman Chemical). Drug pressure was applied 48 h after transfection, using blasticidin (BSD) at a concentration of 2.5 µg/mL, selecting for pKD-VAC-Apt and pKD-GAPM1-mNG-Apt expression. After parasites grew back from transfection, integration was confirmed by PCR, and then cloned using limiting dilution. Clones were maintained in mediums containing 0.5 µM aTc and 2.5 µg/mL BSD.

For generation of VAC^apt^/SBP1^TbID^ parasites, a mix of two plasmids: pTOPO-SBP1-TbID and pUF1-Cas9-SBP1gRNA, was transfected into VAC^apt^ parasites. Drug pressure was applied 48 h after transfection, using 1µM DSM1 ^70^ and selecting for Cas9 expression. After parasites grew back from transfection, integration was confirmed by PCR, and then cloned using limiting dilution.

### Growth assays

For all assays, aliquots of parasite cultures were incubated in 8 µM Hoechst 33342 (ThermoFisher Scientific) for 20 min at room temperature and then fluorescence was measured using a CytoFlex S (Beckman Coulter) flow-cytometer. Flow cytometry data were analyzed using FlowJo software (Tree Star, Inc.) and plotted using Prism (GraphPad Software, Inc.).

For the SBP1^TbID^ growth assay, asynchronous parasites were transferred to a 96-well plate at 0.5% parasitemia and grown for 4 days. Parasitemia was monitored every 24 h.

For the VAC^apt^ and GAPM1^mNG-apt^ growth assays, synchronous ring-stage parasites were washed 5 times with RPMI 1640 medium and split into two cultures, one resuspended in medium containing 0.5 µM aTc and 2.5 µg/mL BSD, and the other one in medium containing only 2.5 µg/mL BSD. Then cultures were transferred to a 96-well plate at 0.2% parasitemia and grown for 6 days. Parasitemia was monitored every 48 h.

### Western blotting

For SBP1^TbID^ parasites, RIPA buffer (150 mM NaCl, 20mM Tris-HCl pH 7.5, 1mM EDTA, 1% SDS, 0.1% Triton X-100) and sonication were used to disrupt parasite pellets conserving all exported proteins. Briefly, late-stage parasites were isolated using a Percoll gradient (Genesee Scientific). Pellets were then resuspended in RIPA buffer and sonicated 3 times at 20% amplitude for 20 s. Protein supernatants were subsequently solubilized in protein loading dye with Beta-mercaptoethanol (LI-COR Biosciences) and used for SDS-PAGE.

For VAC^apt^ and GAPM1^mNG-apt^ parasites, ice-cold 0.04% saponin in 1x PBS was used to isolate parasites from host cells. Parasite pellets were subsequently solubilized in protein loading dye with Beta-mercaptoethanol (LI-COR Biosciences) and used for SDS-PAGE.

Primary antibodies used in this study were mouse-anti-V5 (Cell Signaling Technology, 1:1000), rabbit-anti-PfEF1α (from D. Goldberg, 1:2000), and mouse-anti-HA 6E2 (Cell Signaling Technology, 1:2000). Secondary antibodies used were IRDye 680 CW goat-anti-rabbit IgG, IRDye 800CW goat-anti-mouse IgG, and IRDye 800CW Streptavidin (Li-COR Biosciences, 1:20 000 and 1:10 000). Membranes were imaged using the Odyssey Clx Li-COR infrared imaging system (Li-COR Biosciences). Images were processed and analyzed using ImageStudio (Li-COR Biosciences).

### Immunofluorescence microscopy

For IFAs, cells were fixed as described previously ^67^. The cell lines SBP1^TbID^ and GAPM1^apt^ were smeared on a slide and fixed with acetone. The cell lines VAC^apt^ and VAC^apt^/SBP1^TbID^ were fixed with 4% paraformaldehyde (Electron Microscopy Sciences) and 0.03% glutaraldehyde. Primary antibodies used were mouse-anti-V5 TCM5 (eBioscience, 1:100), rabbit-anti-V5 D3H8Q (Cell Signaling technology, 1:100), rabbit-anti-HA 71550 (ThermoFisher Scientific, 1:100), rabbit-anti-MAHRP (from H. Beck, 1:500), mouse-anti-EXP2 7.7 and mouse-anti-KAHRP (from D. Cavanagh; 1:1000, 1:500 respectively). Secondary antibodies used were Alexa Fluor 488, Alexa Fluor 546, and Streptavidin Alexa Fluor 488 (Life Technologies, 1:1000). Cells were mounted using ProLong Diamond with 4’,6’-diamidino-2-phenylindole (DAPI) (Invitrogen) and imaged using a DeltaVision II microscope system with an Olympus Ix-71 inverted microscope. Images were collected as a Z-stack and deconvolved using SoftWorx, then displayed as a maximum intensity projection. Adjustments to brightness and contrast were made for display purposes using Adobe Photoshop.

### Synchronization assays

For detection of SBP1 during export, SBP1^TbID^ parasites were synchronized with two series of 5% sorbitol treatment. Then, schizont-stage parasites were isolated using Percoll gradient (Genesee Scientific) and immediately transferred to previously warmed fresh red blood cells at 1% hematocrit. Parasites were allowed to egress and invade new red blood cells, and samples were obtained for IFAs at different time points.

### SBP1^TbID^ proximity biotinylation and mass spectrometry

Biotinylation by TbID-tagged SBP1 was confirmed by collecting SBP1^TbID^ parasites for Western blotting and IFAs after incubation for 2 h in biotin-free media supplemented with 50 µM biotin.

For detection of SBP1 during export, SBP1^TbID^ parasites were synchronized with two series of 5% sorbitol treatment. Then, late-schizont-stage parasites were isolated using a Percoll gradient (Genesee Scientific). Parasites were then split in two, and immediately transferred to red blood cells at 1% hematocrit in warm medium without or with biotin (50 µM). Both parasite cultures were incubated for 4 h at 37°C with shaking to let them egress and invade new red blood cells. Cultures were then treated with 5% sorbitol to remove remaining late-stage parasites. Biotinylated culture was washed in 1X PBS, incubated on ice for 10 min to inactivate biotinylation and then stored at -80 °C, until processing. Non-biotinylated culture was incubated for 16 h at 37°C with shaking, then incubated for 4 h in medium with biotin and finally collected as previously described.

Parasite pellets were lysed using extraction buffer (40 mM Tris-HCL pH 7.6, 150 mM KCl, 1mM EDTA, 5% NP-40 and 1X HALT) and sonication (3x, 10% amplitude, 20 s pulses). Streptavidin MagneSphere Paramagnetic Particle beads (Promega) were used to isolate biotinylated proteins. Beads were washed three times in 1 mL of 1X PBS. Protein lysates were incubated with the Streptavidin beads for 1 h at room temperature. After removal of the unbound fraction, the magnetic beads were washed twice with an extraction buffer and once in 1X PBS. The biotinylated proteins on the magnetic beads were digested and analyzed at the Proteomics and Metabolomics shared resource at Fred Hutchinson Cancer Research Center using a Orbitrap Fusion with ETD Mass Spectrometer. All proteins identified in this study are in Supplemental Table 2. All mass spectrometry proteomic data have been deposited to the ProteomeXchange consortium via the MassIVE partner repository with the dataset identified PXD034946 (Project name: Rapid proximity biotinylation of the *Plasmodium falciparum* exported protein, SBP1).

## SUPPLEMENTAL FIGURES

**Supplemental figure 1.**
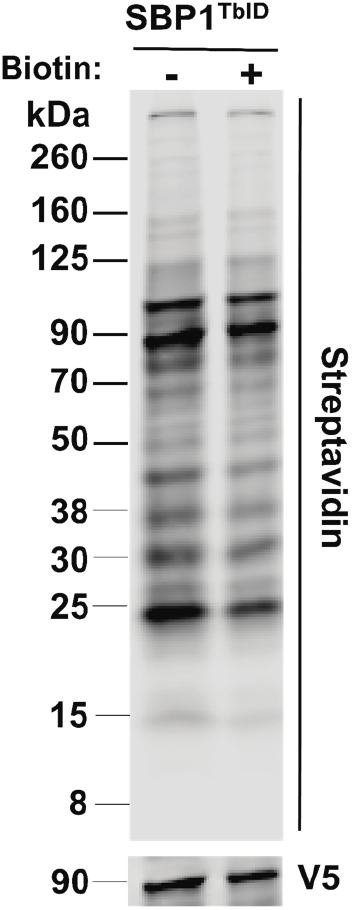
Biotinylation of proximal proteins by TurboID_V5_-tagged SBP1. Western blot of parasite lysates isolated from the mutant line SBP1^TbID^ grown in complete RPMI medium, incubated with or without biotin (50 μM) for 2 h. Samples were probed with antibodies against V5 (loading control) and fluorescent dye-labeled streptavidin. The protein marker sizes are shown on the left.

**Supplemental figure 2.**
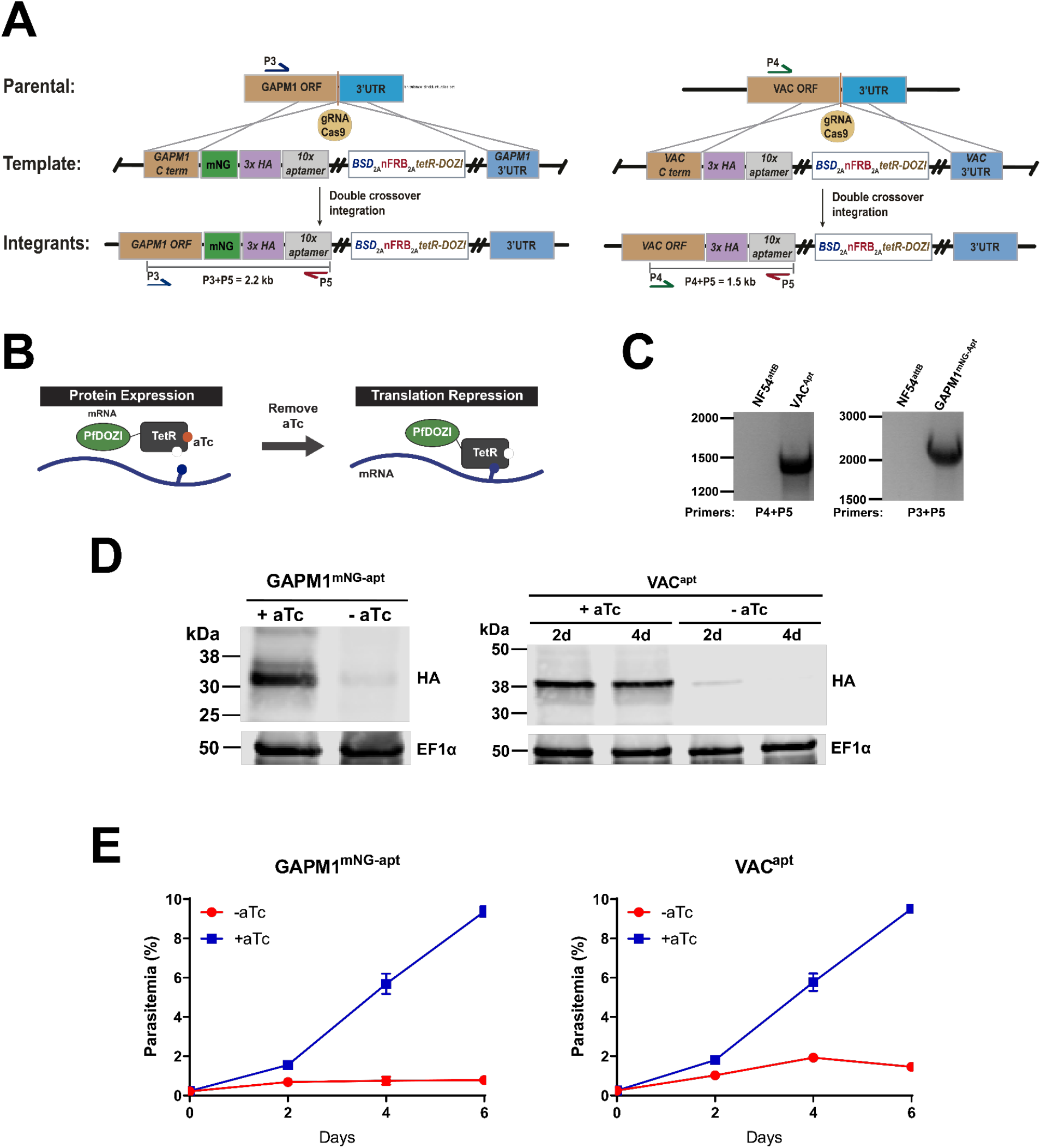
Characterization of parasite lines VAC^apt^ and GAPM1^mNG-apt^. (A) Schematic showing the integration of the repair plasmid to modify the genomic loci of Pf3D7_1432100 (VAC) and Pf3D7_1323700 (GAPM1). Cas9 introduces a double-stranded break at the C-terminus of the VAC and GAPM1 locus. The repair plasmid provides homology regions for double-crossover homologous recombination, introducing the HA-tag and the TetR-Aptamer system. For GAPM1^mNG-apt^, a fluorescent tag mNeonGreen was introduced between the C-terminus and the HA-tag. (B) Regulation of protein expression using the TetR-Aptamer knockdown system. TetR binds to aptamer repeats in the mRNA, while PfDOZI localizes the complex to sites of mRNA sequestration, causing a repression in translation of the gene of interest. Anhydrous tetracycline (aTc) binds to TetR, blocking its interaction with the aptamers. (C) PCR test confirming integration at the VAC and GAPM1 locus. Amplicons were amplified from genomic DNA isolated from mutant and wild-type parasites. Primers were designed to amplify the region between the C-terminus and the tandem of 10X aptamer repeats. (D) Western blot of parasite lysates isolated from the mutant lines VAC^apt^ and GAPM1^mNG-apt^ probed with antibodies against HA and EF1α (loading control). The protein marker sizes are shown on the left. GAPM1^mNG-apt^ parasites were collected after incubation for 48 h in the presence or absence of aTc. VAC^apt^ parasites were collected after incubation for 48 and 96 h in presence or absence of aTc. (E) Growth of synchronous VAC^apt^ and GAPM1^mNG-apt^ parasites over 6 days after removal of aTc from the medium via flow cytometry. Representative of three biological replicates shown for each growth curve. Each data point represents the mean of three technical replicates; error bars represent standard deviation.

**Supplemental figure 3.**
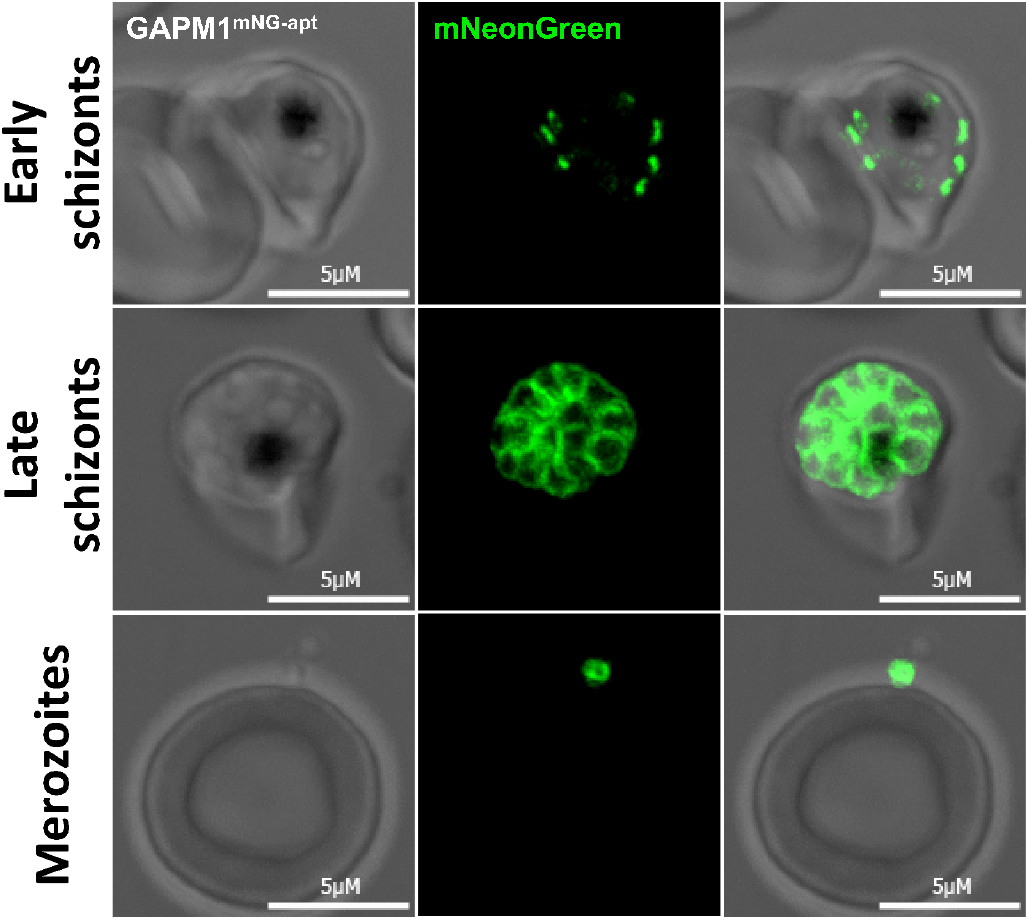
Localization of GAPM1 at late stages in the GAPM1^mNG-apt^ cell line. (A) Representative live images showing GAPM1^mNG-apt^ localization at early and late schizonts, and merozoites. Images of GAPM1^mNG-apt^ from left to right are phase-contrast, mNeonGreen (green), and fluorescence merge. Z stack images were deconvolved and projected as a combined single image.

**Supplemental figure 4.**
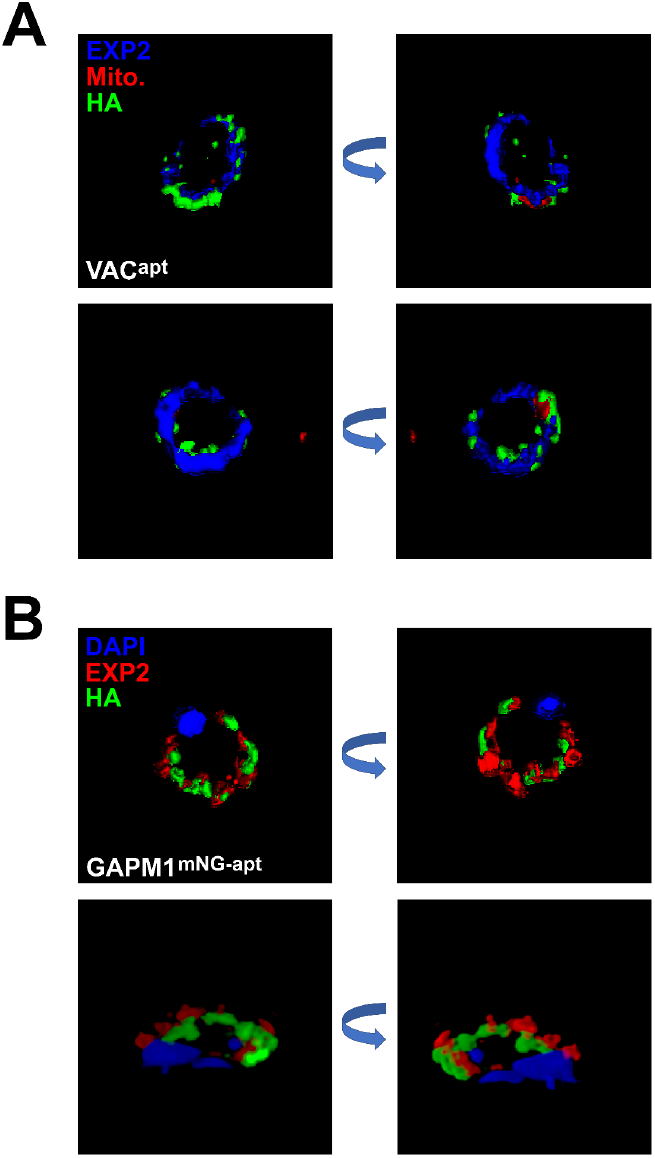
Localization of VAC and GAPM1 at early stage parasites. 3D reconstruction based on structured illumination microscopy images captured from (A) VAC^apt^ and (B) GAPM1^mNG-apt^ ring-stage parasites at 4 hpi and stained with the antibodies as in Fig 5A.

**Supplemental Table 1**. List of primers used in the study to generate the cell lines SBP1^TbID^, VAC^apt^ and GAPM1^mNG-apt^

**Supplemental Table 2**. Complete list of proteins identified using label-free analysis and collected by mass spectrometry.

