## Supplemental Table 1 for "Rapid, time-resolved proximity labeling by sbp1 identifies a porin domain protein at the malaria parasite periphery"

| Primer | Amplicon | Sequence (5’ – 3’) |
| --- | --- | --- |
| P1 | SBP1-Cterm | CCCTCACTAAAGGGACTAGTCTTTGTTATTAACATATTATTGTTCATCAACTTTTACAAC |
| P2 | SBP1-Cterm | TAGATCTGTTAACGGATCCGGTTTCTCTAGCAACTGTTTTTGTCGTTGATTTGGTTGTGG |
| P3 | SBP1-3’UTR | TGGACAGCACCTAAGAATTCAGATAAATTATTATAAATCAATTGTGCCAACAATAATGAG |
| P4 | SBP1-3’UTR | TAGCGGCCGCGAATTCGTTGTGAACGTTTTTAATTATGTATGCATACAAAAAATATAC |
| P5 | TurboID | CCCTCACTAAAGGGACTAGTGCTCGGGATCCACCGGTCGCCACCATG |
| P6 | TurboID | TAGCGGCCGCGAATTCTTAGGTGCTGTCCAGGCCCAGCAGGGGGTTG |
| P7 | VAC-Cterm | TGCAGAAAGGTGTGGATATCATCCCGAGTAATAAACACTTTTATGGATCC |
| P8 | VAC-Cterm | CGTCATAAGGGTATCCGGAGACGTCTGATTTTAAATAAAGTTTCATTCCAAATTTGGTG |
| P9 | VAC-3’UTR | TCCAATGGCCCCTTTCCGGGCGCGCCTCTTATTTGTTTTTATTTATTAAGGAAGAATTAG |
| P10 | VAC-3’UTR | TTATTACTCGGGATGATATCCACACCTTTCTGCACCTTATATATAC |
| P11 | GAPM1-Cterm | TGGTGCTAGGTAGGGATATCGAACTGTATCATGGAGCTTGTCCCTTATATGTTTG |
| P12 | GAPM1-Cterm | AAACGGTGGCGACCGGTGGATCCCGAGCACATTGTTTGCATGCTGCAATATTTTCGGTAG |
| P13 | GAPM1-3’UTR | TCCAATGGCCCCTTTCCGGGCGCGCCCCTACAAATTAACAAATTCGAAGAATACAAAAG |
| P14 | GAPM1-3’UTR | CCATGATACAGTTCGATATCCCTACCTAGCACCACATTTTAACATTG |
| P15 | mNeonGreen | ATGTGCTCGGGATCCACCGGTCGCCACCGTTTCTAAGGGTGAAGAAGATAACATGGCTTC |
| P16 | mNeonGreen | CGTCATAAGGGTATCCGGAGACGTCCTTGTATAATTCATCCATACCCATAACATCAGTG |
| P17 | SBP1 gRNA | TAAGTATATAATATTTCTAGCAACTGTTTTTGTTGGTTTTAGAGCTAGAA |
| P18 | SBP1 gRNA | TTCTAGCTCTAAAACCAACAAAAACAGTTGCTAGAAATATTATATACTTA |
| P19 | VAC gRNA | CATATTAAGTATATAATATTTACTGTCTATAATTAAACAAGTTTTAGAGCTAGAAATAGC |
| P20 | GAPM1 gRNA | CATATTAAGTATATAATATTATGCAAACAATGTTAAAAAGGTTTTAGAGCTAGAAATAGC |
